# On the relationship between high-order linkage disequilibrium and epistasis

**DOI:** 10.1101/305342

**Authors:** Yanjun Zan, Simon K. G. Forsberg, Örjan Carlborg

## Abstract

A plausible explanation for statistical epistasis revealed in genome wide association analyses is the presence of high order linkage disequilibrium (LD) between the genotyped markers tested for interactions and unobserved functional polymorphisms. Based on findings in experimental data, it has been suggested that high order LD might be a common explanation for statistical epistasis inferred between local polymorphisms in the same genomic region. Here, we empirically evaluate how prevalent high order LD is between local, as well as distal, polymorphisms in the genome. This could provide insights into whether we should account for this when interpreting results from genome wide scans for statistical epistasis. An extensive and strong genome wide high order LD was revealed between pairs of markers on the high density 250k SNP-chip and individual markers revealed by whole genome sequencing in the *A. thaliana* 1001-genomes collection. The high order LD was found to be more prevalent in smaller populations, but present also in samples including several hundred individuals. An empirical example illustrates that high order LD might be an even greater challenge in cases when the genetic architecture is more complex than the common assumption of bi-allelic loci. The example shows how significant statistical epistasis is detected for a pair of markers in high order LD with a complex multi allelic locus. Overall, our study illustrates the importance of considering also other explanations than functional genetic interactions when genome wide statistical epistasis is detected, in particular when the results are obtained in small populations of inbred individuals.

## Introduction

The genetic architecture of most biological traits is complex and involves multiple genes, whose effects are often influenced by interactions with other genes and environmental factors. To study the relative contributions by genes, environmental factors and their interactions in segregating populations, statistical genetic approaches are commonly used to partition the genetic variance to additive and dominance variance of individual loci and epistatic interaction variance between them (Lynch and Walsh 1998). In principle, the variance partitioning is performed by associating the phenotypic variation for a trait in a population with linear combinations of the genotypes within and/or across loci. How the genotypes are combined (parameterized) in the model is determined by the genetic model used in the analysis. The classic quantitative genetics models are parameterized to capture the genetic variance in a hierarchical manner. First, a main additive allele-substitution is defined. Then, if accounted for, dominance is modeled as a single-locus deviation from additivity and genetic interactions as multi-locus deviations from single locus additivity and dominance (Nelson *et al.* 2013). As a consequence of this, the genetic contributions of individual and combinations of loci described as additive, dominance and epistatic variances are unlikely to reflect the underlying biological mechanisms (Carlborg *et al.* 2006; Phillips 2008; Huang *et al.* 2012; Sackton and Hartl 2016; Forsberg *et al.* 2017).

Although the ultimate aim of a genetic association study is generally to detect functional polymorphisms, most often genotypes are only scored for a reduced set of polymorphisms (genetic markers). These reduced marker sets are selected with the aim to tag as many of the unobserved functional polymorphisms as possible. The statistical inferences of the underlying genetic architecture made from such reduced sets of markers can, however, be problematic in some cases. For example, multiple unobserved functional polymorphisms can lead to associations to individual markers that do not properly represent the causal variants (Platt *et al.* 2010), and high order linkage disequilibrium (LD) to single functional polymorphism can lead to indirect statistical epistatic associations to pairs of markers (Wood *et al.* 2014). Here, we focus on high order linkage disequilibrium defined as when two genotyped markers tag an un-genotyped polymorphism (see Materials and Methods section). It is still unknown how prevalent and strong such high order LD is in the genome, making it difficult to estimate how many reported pairwise statistical epistatic interactions are due to such LD. However, the study by *Wood et al* (Wood *et al.* 2014) presented results suggesting that many of the significant statistical epistatic interactions detected between pairs of local markers by Hemani *et al.* (Hemani *et al.* 2014) might be due to high-order LD to unobserved, linked sequence polymorphisms in the same genomic region. Many past and current studies of genetic interactions in, for example, Drosophila, plant, animal and human populations (Shimomura *et al.* 2001; Anholt *et al.* 2003; Caicedo *et al.* 2004; Segrè *et al.* 2004; Carlborg *et al.* 2006; Hemani *et al.* 2014) rely on genome-wide statistical analyses of pairwise interactions between selected sets of markers as in (Hemani *et al.* 2014). With the increasing interest in, and availability of, sufficiently large datasets for epistatic association analyses it is therefore important to also evaluate the risk of making false inferences about loci being involved in functional genetic interactions from findings of statistical epistasis, when they instead are due to high order LD.

Here, we empirically explore the prevalence and strength of high order LD within and between chromosomes in publically available high-density SNP and whole-genome re-sequencing data from the model plant *Arabidopsis thaliana.* Two locus LDs are calculated between the markers selected for the 250k *A. thaliana* SNP chip that have been the basis for many GWAS analyses in the past, and the additional SNPs revealed by whole genome sequencing using data from the 1001 genomes project (Atwell *et al.* 2010; Cao *et al.* 2011; Horton *et al.* 2012; Schmitz *et al.* 2013; Alonso-Blanco *et al.* 2016). Strong high order LD was found to be common both within and across chromosomes between pairs of markers from the SNP-chip and the sequencing polymorphisms and often the combined genotype of the marker pair tagged the genotype of the sequencing markers better than any single marker on the SNP chip. The risk of falsely inferring genetic interactions between markers on different chromosomes in a two-locus interaction analysis might increase in situations when the underlying genetic architecture is more complex, for example when a single locus contains multiple functional alleles. This is illustrated using an empirical example from a second public *A. thaliana* dataset (Forsberg *et al.* 2015). Overall, this study provides new insights that deepen our understanding about the link between high order LD and statistical epistasis to guide researchers when interpreting results obtained from epistatic genetic association analyses.

## Materials and Methods

### Methods

When an individual marker is in complete linkage disequilibrium (r^2^ = 1) with a functional polymorphism affecting a studied trait, a single-locus association test between the marker and the trait will capture all the phenotypic variance contributed by the functional polymorphism. A basic assumption in genetic association studies is that at least one genotyped marker will be in sufficiently high LD with each functional polymorphism to detect it in this way. In reality, however, not all functional polymorphism will be in such perfect LD with a genotyped marker, and then there is a risk that the joint genotype of two (or more) markers tags the genotype of the functional polymorphism better than any single marker (high-order LD > single-marker LD). This will, as discussed below, influence the significances of the trait-marker associations detected in a genetic association analysis and the inferences made about the genetic architecture of the trait.

#### Quantifying high order linkage disequilibrium

We calculate the high order LD between pairs of predictors (here genotyped SNP markers) and single targets (here un-genotyped SNP polymorphisms) following (Hao *et al.* 2007).

Consider a pair of bi-allelic predictor SNPs (M_1_ and M_2_; Figure 1). These markers can together form four two-locus genotypes: AB, Ab, aB and ab (Figure 1). We now want to know whether any two-locus predictor could tag the single locus target genotype better than any of the individual predictor genotypes (i.e. evaluate whether max(second-order LD) > max(single order LD)). To calculate the high order LD between the two predictors (M_1_ and M_2_) and the single target (Q), the two-locus M_1_M_2_ genotype is used to create a multi-allelic pseudo marker (P) with four alleles (Figure 1). In this way, a second-order LD (r^2^) can be calculated for each of the possible ways that M1 and M2 together can tag the genotype at Q (Figure 1).

**Figure 1.**
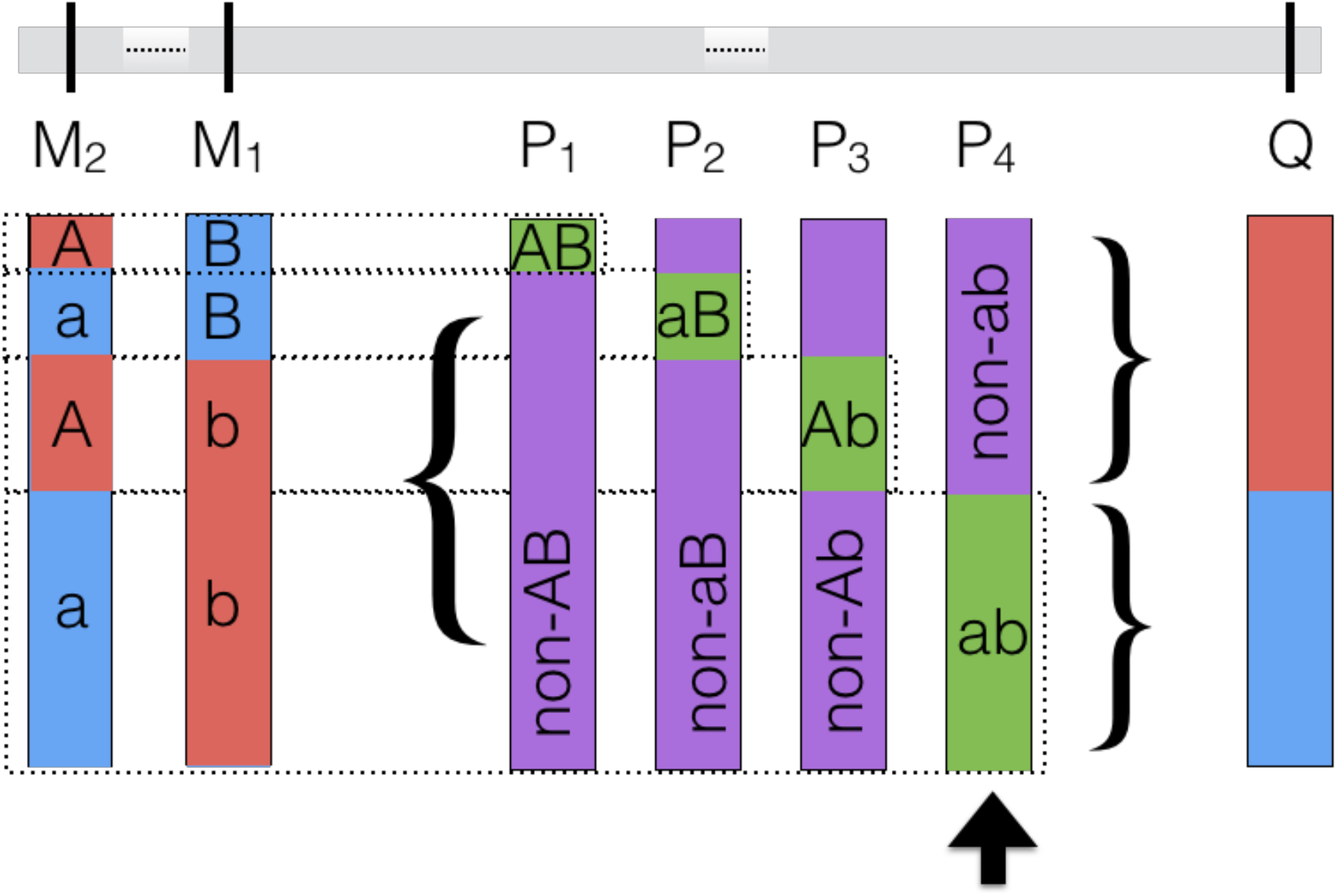
Illustration of how the pseudomarkers (P_i_, P_2_, P_3_, P_4_) used in the estimation of the second order linkage disequilibrium between a pair of linked or unlinked markers (predictors; M_1_ and M_2_,) and a third linked or unlinked functional polymorphism (target; Q) are created. The pseudomarkers together represent the possible bi-allelic formulations of the two-locus M_1_M_*2*_ genotypes. The maximum pairwise LD-r^*2*^ between the target and the four pseudomarkers (P_4_) defines the second order LD between the predictors (M_1_, M_2_) and the target (Q).

The calculation of the second order LD therefore first involves creating the four possible bi-allelic pseudomarkers (P_1_, P_2_, P_3_ & P_4_; Figure 1) from the two locus M_1_/M_2_ genotypes. These are assigned the genotypes P_1_{AB, non-AB}, P_2_{Ab, nonAb}, P_3_{aB, non-aB} and P_4_{ab, non-ab}, respectively. The LD-r^2^ is then computed between the target (Q) and the four bi-allelic pseudomarkers (P_1_, P_2_, P_3_ & P_4_). For each pair of predictors, the second order LD is then defined as the LD-r^2^ for the pseudomarker with the highest LD-r^2^ to the target. Pseudomarkers with higher LD-r^2^ to the target (Q) than 0.3 are kept for further analyses. The LD-r^2^ values were computed using the software *LdCompare* (Hao *et al.* 2007).

#### Statistical epistasis emerging from high order linkage disequilibrium

In a genetic association study in an inbred or haploid population, two-locus epistasis is typically modelled as:

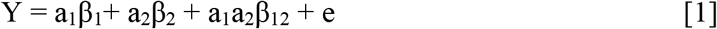

Here, a_1_ and a_2_ are indicator variables for the genotypes at two genotyped markers, M_1_ and M_2_, taking values 1/−1 for the two alternative homozygous genotypes AA vs aa and BB vs bb, respectively. a_1_a_2_ is an indicator variable for the interaction between a_1_ and a2 taking value 1 for the two-locus genotypes AABB and aabb and −1 for AAbb and aaBB. β_1_, β_2_ and β_12_ are the corresponding estimates for the marginal (additive) effects and the additive-by-additive interaction between the loci.

The aim of a statistical epistatic analysis is to include an interaction term in the model [1] to estimate the deviations of the two-locus genotype-values (AABB, AAbb, aaBB and aabb) from the predictions obtained by the marginal (additive) effects (Alvarez-Castro and Carlborg 2007). However, a non-zero estimate of the interaction term in model [1] does not, as noted e.g. by Wood et al. (Wood *et al.* 2014) necessarily have to result from a genetic interaction. It could, for example, instead emerge from a second-order LD between two markers and a single functional polymorphism. Here, refer back to Figure 1. Now assume that a trait is determined by a single functional locus (Q). Two markers, M_1_ and M_2_, are genotyped but neither of these markers individually tag the causal genotype (blue) at Q well. However, the causal (blue) allele at Q is,tagged perfectly by one of the two-locus M_1_M_2_ genotypes (ab; Figure 1), while the other three M_1_M_2_ two-locus genotypes (aB, Ab and AA; Figure 1) are only present together with the no-effect (red) allele at locus Q. When fitting model [1] to the genotypes of marker M_1_ and M_2_, the estimate for the interaction term (β_12_) will be non-zero, illustrating how statistical epistasis can emerge from the second-order LD between M_1_ and M_2_ and Q. This example illustrates a scenario similar to what was empirically observed in (Wood *et al.* 2014), where physically linked markers in low LD with each other tagged haplotypes that were in high order LD with a polymorphism that was unobserved in the original study.

#### Classifying identified high order linkage disequilibrium triplets depending on the distance between the loci

Here, we evaluate the prevalence and strength of high order LD between pairs of markers selected for genotyping on a 250k SNP chip (predictors) and a third locus revealed by whole genome sequencing (targets) using publicly available datasets in *A. thaliana* (Cao *et al.* 2011; Alonso-Blanco *et al.* 2016). Three types of high order LD are defined based on the locations of the predictors relative to the target. If both predictors are located within 1Mb of the target it is classified as cis-cis. If only one predictor is closer than 1Mb it is classified as cis-trans. If none is closer than 1Mb it is classified as trans-trans. The choice of a 1Mb threshold to define cis vs trans predictors is arbitrary, but we consider it useful for evaluating how common high order LD is between predictors near (local/cis) and far (global/trans) from the target.

### Material

#### The genome wide prevalence of high order linkage disequilibrium in publically available *Arabidopsis thaliana* datasets

The *A. thaliana* 1001-genomes project has released complete genome sequences for hundreds of wild collected accessions (http://www.1001genomes.org). Here, we used whole-genome SNP data on 728 accessions scored by whole genome re-sequencing (Cao *et al.* 2011; Alonso-Blanco *et al.* 2016). The predictors used in our analysis was a subset of the SNPs selected for the 250k *A. thaliana* SNP chip (Horton *et al.* 2012) (n = 200,352 in total; MAF > 0.05) and the targets a subset of the SNPs revealed using whole-genome re-sequencing (n = 1,641,240 in total; MAF > 0.05) (Table 1). Although the results from the analyses of this data will be specific to this species and dataset, it is assumed that the relationships between targets and predictors will be a realistic representation of what to be expected also in other populations. This is because the selection of markers for the high-density 250k SNP chip, was done for the purpose of genetic association studies following similar procedures as used also in other species and populations.

**Table 1.** Regions and SNPs selected for evaluation of second order LD.

|  | Window 1 | Window 2 | Targets<sup>1</sup> | Predictors<sup>2</sup> | Filtered targets<sup>3</sup> |
| --- | --- | --- | --- | --- | --- |
| Region 1 | Chr2: 8-14Mb | - | 70,712 | 6,053 | - |
| Regionpair 1 | Chr1: 10-12Mb | Chr3: 10-12Mb | 29,133 | 6,245 | 20,239 |
| Regionpair 2 | Chr2: 10-12Mb | Chr4: 10-12Mb | 23,751 | 5,302 | 15,887 |
| Regionpair 3 | Chr2: 10-12Mb | Chr3: 10-12Mb | 23,751 | 5,212 | 15,884 |
| Genome |  |  | 1,486,942 | 154,298 | 1,229,012 |

^1^Total number of polymorphic SNPs in the evaluated windows/genome in the population revealed via whole-genome re-sequencing (Alonso-Blanco *et al.* 2016). ^2^Total number of polymorphic SNPs in the two windows/genome included on the 250k AT SNP-chip (*Horton et al.* 2012); ^3^Number of target SNPs in the two windows/genome with LD-r^*2*^ < 0.6 to any individual predictor.

The reason for only studying a subset of the possible targets and predictors is that it was not computationally feasible to exhaustively evaluate the high order LD between all possible pairs of predictors selected for the 250k SNP chip and all the targets revealed by genome sequencing. Instead, the second order LD was exhaustively calculated for all targets and predictors i) within a randomly selected 6 Mb window on chromosome 2 as well as ii) between three randomly selected windows from different chromosomes (Table 1). Computations were performed for the entire population (n = 728 individuals) and two smaller random samples of n = 100 and n = 50 individuals. The results for the populations with n = 100 and n = 728 are reported in the main manuscript and the results for n = 50 is reported in the Supplementary material.

The predictor pairs in the evaluated windows in the genome with high order LD-r ^2^ > 0.6 to a target were classified as cis-cis/cis-trans/trans-trans. To extrapolate these findings to the genome level, the proportions of all evaluated predictor pairs that displayed these patterns were calculated and then multiplied with the total number of possible cis-cis/cis-trans/trans-trans pairs in the genome (Table S1).

#### *Analyzing a public* A. thaliana *dataset for two locus statistical epistasis*

A publicly available dataset including 340 *Arabidopsis thaliana* accessions were used for a genome wide association analysis. In short, the plants were grown in a controlled environment with 6 biological replicate plants per accession. Analyses by Inductively Coupled Mass Spectroscopy (ICP-MS) provided estimates of leaf molybdenum concentration as described in (Baxter *et al.* 2010; Forsberg *et al.* 2015). The accessions were genotyped for 141,385 SNP markers with MAF > 0.15 (Atwell *et al.* 2010; Baxter *et al.* 2010; Shen *et al.* 2012; Forsberg *et al.* 2015). A more thorough description of the dataset can be found in (Baxter *et al.* 2010; Forsberg *et al.* 2015). In an earlier study of this dataset (Forsberg *et al.* 2015), it was revealed that a large fraction of the genetic variance for this trait was explained by a single linkage block containing several low-frequency, large effect structural variants that were poorly tagged by the genotyped SNPs. This linkage block was originally identified due to its large marginal, variance heterogeneity effect in the population (Shen *et al.* 2012). It is known that statistical epistasis and genetic variance heterogeneity can emerge from similar genetic architectures (Forsberg *et al.* 2015), and this population was therefore selected for further evaluations of whether high order LD between the genotyped SNPs and these hidden polymorphisms could lead to statistical epistasis in a two locus association analysis. We performed an exhaustive, two-dimensional genome scan for pairwise statistical epistasis between the genotyped markers and the level of molybdenum in the leaf using the software *plink* (Purcell *et al.* 2007) without control for population structure. Thereafter, each pair of loci that passed the genome wide significance threshold in the initial scan was fitted in a two-locus epistatic genetic model [1] using *hglm* function in *hglm* package (Rönnegård *et al.* 2010) to correct for the possible effects of population structure via the genomic kinship matrix as in (Forsberg *et al.* 2015). The significance threshold used to infer significant interacting pairs (p < 3.2 × 10^−10^) was defined as a Bonferroni corrected nominal 5% significance threshold. The correction was done for an estimated number of independent association tests assumed to equal the number of independent LD blocks in the *A. thaliana* genome as described in (Lachowiec *et al.* 2015).

#### Data availability

Genome wide re-sequencing data are available as part of the *Arabidopsis thaliana* 1001 genomes project http://1001genomes.org/data-center.html. The 250 K SNP chip data are available as part of the genotype data for the *Arabidopsis thaliana* Regmap panel (http://bergelson.uchicago.edu/7pageid=790). The Molybdenum levels for the 340 *Arabidopsis thaliana* accessions are available in https://doi.org/10.1371/journal.pgen.1005648.s005

## Results

This study aims to answer the following questions by analyzing two public *A. thaliana* datasets: How common can we expect high order LD to be between pairs of SNPs selected for genotyping and hidden sequence variants in the genome? Is high order LD primarily observed between predictors tightly linked to a target functional polymorphism (in cis) as in (Wood *et al.* 2014), or is it also observed for predictors unlinked to the target (in trans)? How dependent is the prevalence of high order LD and cis vs trans predictors on the population size? We also present an empirical example where high order LD exists between a cis-trans predictor pair with significant statistical epistasis and a locus displaying a strong genetic variance heterogeneity due to independent contributions by multiple linked polymorphisms (Forsberg *et al.* 2015). This illustrates how complex inheritance patterns of individual loci, something usually not explored in GWAS data, further complicates the interpretation of detected statistical epistatic signals.

### The population size affects the prevalence and location of predictors in high order LD

The high order LD-r^2^ values for all pairs of predictors and individual targets in a 6Mb window on Chromosome 2 (Table 1) is shown for populations with n = 100 and n = 728 individuals in Figure 2. The strongest second order LD-r^2^ was observed where at least one predictor is located near the target (y-axis). When the sample size was smaller (n = 100; Figure 2A), strong second order LD-r^2^ was rather common also when both predictors were located far from the target. For example, 20% of the targets had a high order LD-r ^2^ > 0.65 with a predictor pair where at least one of the predictors was located more than 1Mb away from it. Even though the prevalence of strong high order LD-r^2^ decreases when the sample size increases, it is still common in the large population (n = 728; Figure 2B), with the highest prevalence when at least one of the predictors is located close to the target.

**Figure 2.**
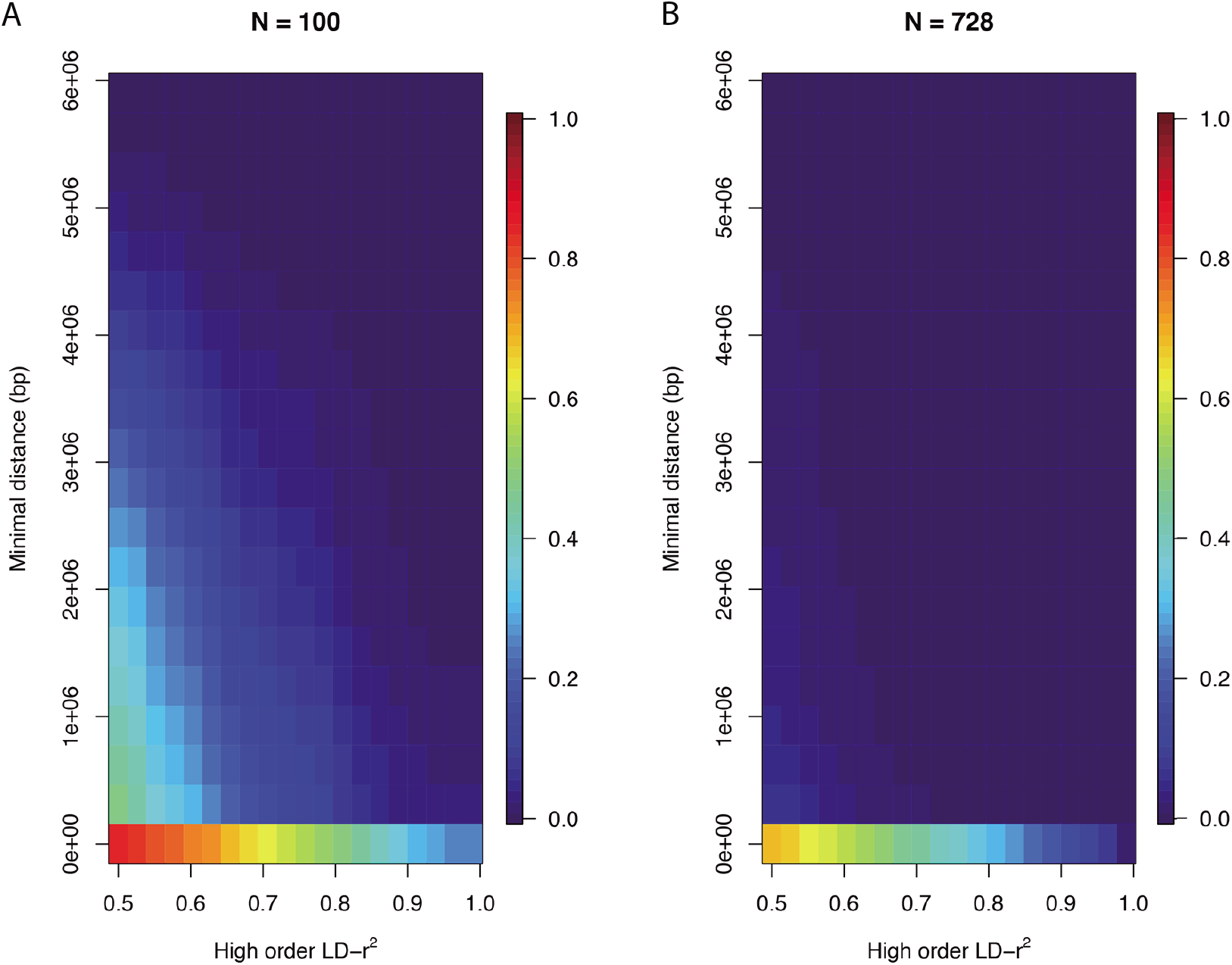
*Illustration of how the prevalence of high order LD-r^2^ to the targets in a 6Mb window on* A. thaliana *chromosome 2 (8 - 14Mb) depends on distance of the predictors from the target. The color gradient illustrates the proportion of predictor pairs that reach a particular LD-r^2^ (x-axis) depending on the distance between the nearest predictor and the target (y-axis). Results are presented for populations with n = 100 (**A**) and n = 728 (**B**) individuals*.

Strong high-order LD-r^2^ between a predictor pair and a target is mostly observed when at least one of the predictors is in strong individual LD-r^2^ with the target. However, as illustrated in Figure 3, many cases also exist where the high order LD-r^2^ is strong while the LD-r^2^ to the individual predictors is weak.

**Figure 3.**
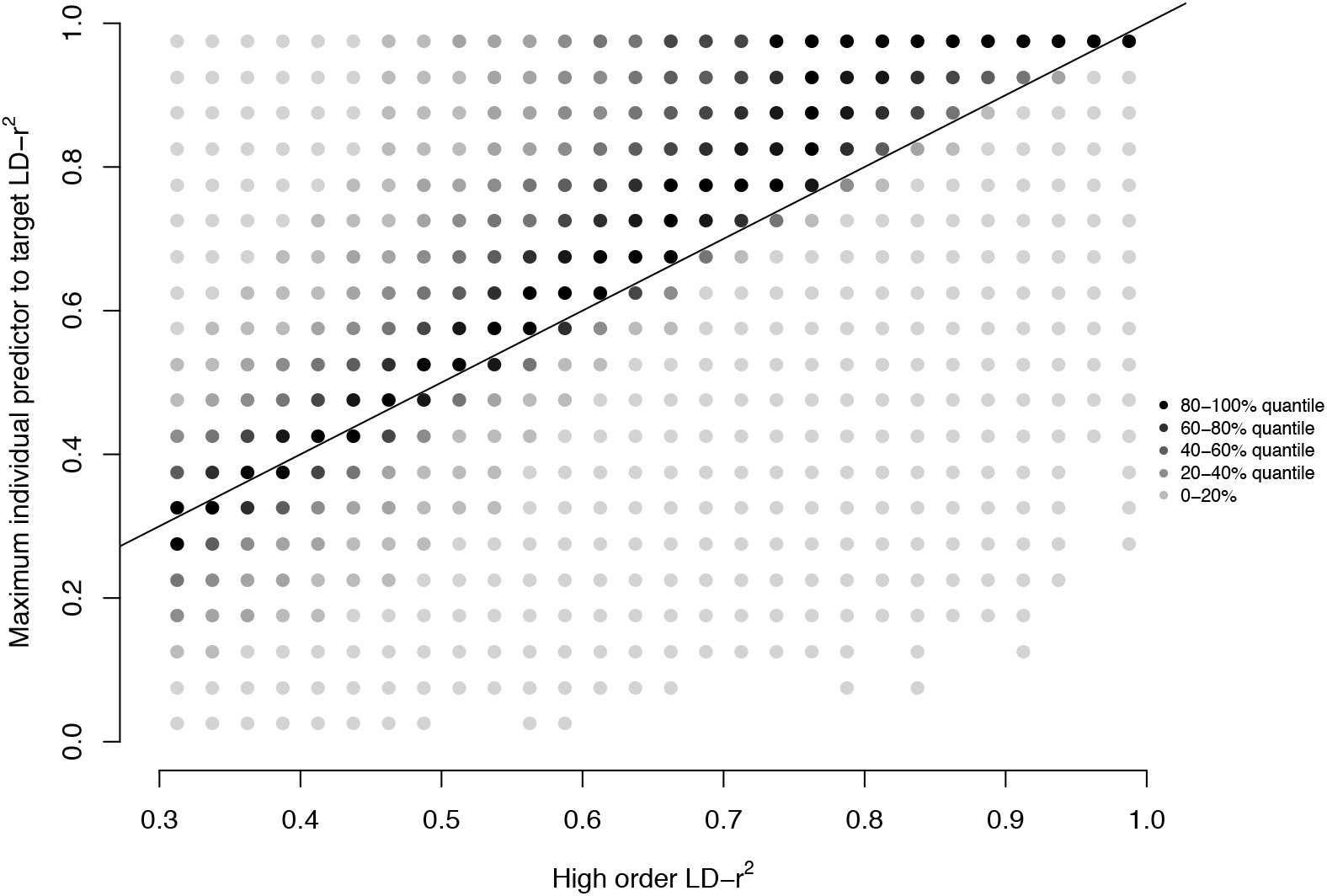
Strong second order LD-r^2^ exists also when the individual predictor to target LD-r^2^ is weak. The intensity of each dot illustrates the number of cases with a particular high order LD-r^2^ / maximum individual predictor to target LD-r^2^ combination. Dots below the line are cases where the high order LD-r^2^ stronger than any individual predictor to target LD-r^2^ (n=728).

### Estimating the genome wide prevalence of strong high order linkage disequilibrium

Figure 2 illustrates that high-order LD-r^2^ exists where one or both predictors are located close to the target as well as when one or both predictors are located further away in the evaluated 6Mb window. The genome-wide prevalence of high order LD-r^2^ for the three different classes of predictor pairs, cis-cis/cis-trans/trans-trans (as defined above) were next explored in three pairs of distant 2Mb windows in the genome (Table 1) to provide data to estimate their genome-wide prevalence. Here, only cases when individual predictors in the windows had lower individual LD-r^2^ than 0.6 to the targets were considered.

Overall, the fraction of predictor pairs that display higher second-order LD (LD-r^2^ > 0.6) is low. In the smaller population (n = 100), less than 1 out of 10^6^ evaluated predictor pairs and in the larger population (n = 768) less than 1 out of 10^7^ (Table S1). However, since the total number of evaluated pairs was very large (around 10^11^), many cases were still detected. Regardless of population size, cis-cis and cis-trans pairs dominated (42/44% for n = 100, and 56/58% for n = 728; Figure 4A-C; Table S1). Trans-trans pairs existed, but were much less common (~1% for n = 100, <0.01% for n = 728, respectively, Figure 4A-C; Table S1). When extrapolating these results to a genome wide scale, this picture, however, changes dramatically (Figure 4D). Trans-trans and cis-trans predictor pairs are now much more common than cis-cis pairs due to their much higher genome-wide prevalence (35/18-fold for n = 100 and 35/0.3 for n = 728 more common; Figure 4D, Table S1). This result illustrates that it is a considerable risk to disregard high-order LD as a possible explanation for statistical epistatic interactions even at larger sample-sizes.

**Figure 4.**
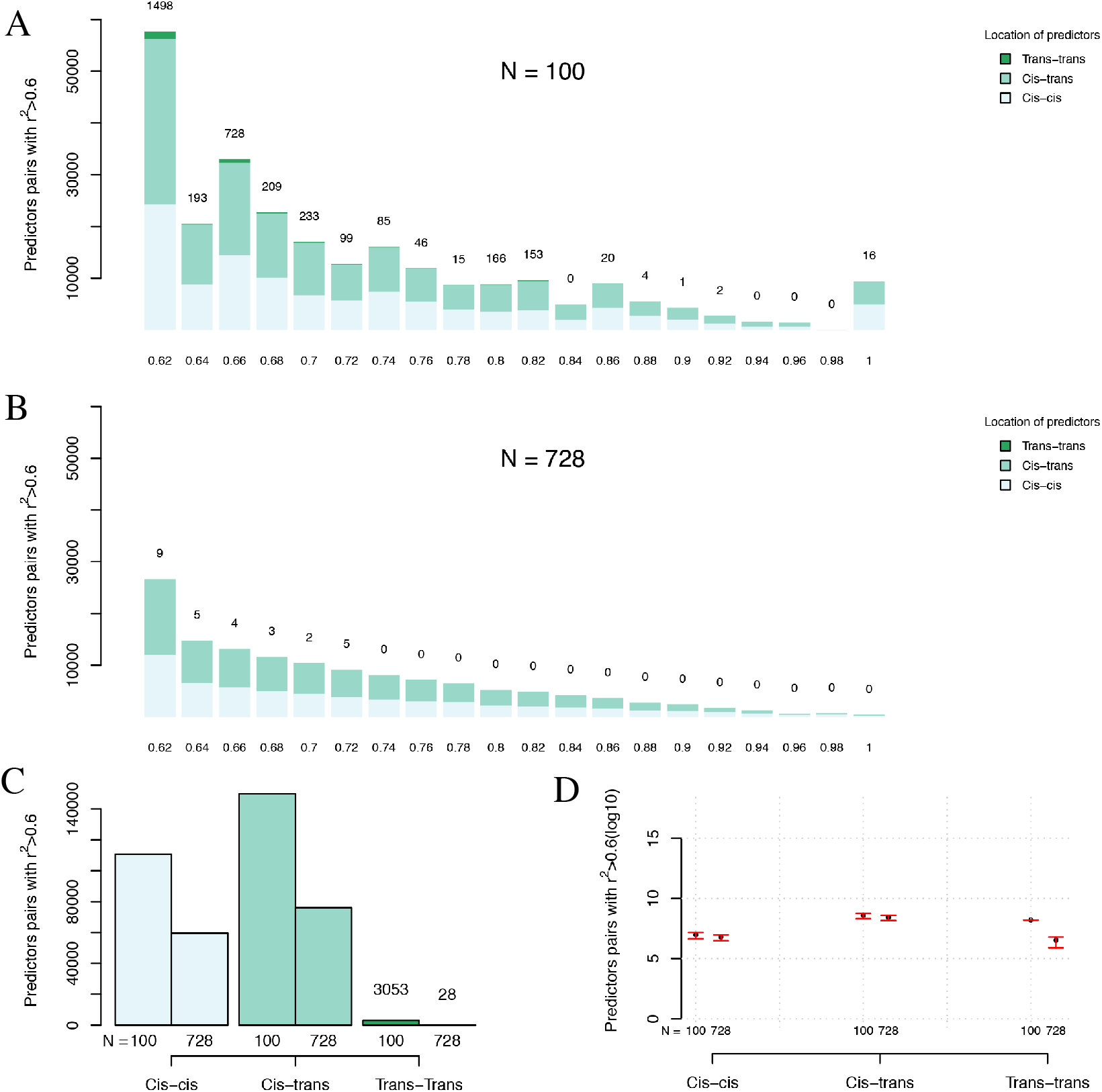
Number of predictor pairs of different classes in strong high order LD-r^2^ (>0.6) to targets detected in the evaluated windows and estimated genome wide. The distribution of LD-r^2^ values > 0.6 for the cis-cis, cis-trans,trans-trans predictor pairs for (**A**; n = 100) and (**B**; n = 728) The total number of predictor pairs with high order LD-r^2^ above 0.6 in the three classes are summarized in (**C**) and used to estimate the total expected number of predictor pairs in the entire genome (**D**; error bars show the estimation error estimated from the results obtained for the three window (Materials and Methods).

### *Linking high order LD and statistical epistasis in a two locus epistatic association analysis in* A. thaliana

A publicly available dataset including 340 *Arabidopsis thaliana* accessions were used for a genome wide association analysis for leaf molybdenum concentration This dataset was earlier used by (Forsberg *et al.* 2015) to dissect a locus with a highly significant variance heterogeneity association for leaf molybdenum concentration (Shen *et al.* 2012) to the contributions of four independent associations in an extended LD block on chromosome 2. Several of these associations were found to structural variants that were poorly tagged by the SNP markers (Forsberg *et al.* 2015). Our pairwise genome wide scans for pairs of epistatic loci identified 396 significant SNP pairs. For 290 pairs both markers were located in the narrow region on chromosome 2 that was earlier dissected in detail (Forsberg *et al.* 2015). All these are examples of cis-cis predictor pairs. The remaining 106 pairs contained one predictor in the chromosome 2 region and another one elsewhere in the genome, being examples of cis-trans predictor pairs.

The strongest pairwise epistasis was detected for a cis-trans predictor pair (Figure 5A). The accessions with the AA genotype at the predictor located in trans to the chromosome 2 region (chromosome 1:5,315,502 bp) all have an intermediate molybdenum level in the leaf (Figure 5A). The accessions with the GG allele at the trans predictor have different levels of molybdenum in their leaves depending on whether they carry the CC or TT genotype at the cis predictor in on chromosome 2 (10,928,720 bp). These differences explain the significant statistical epistasis detected when fitting the two-locus epistatic model [1] to this data.

**Figure 5.**
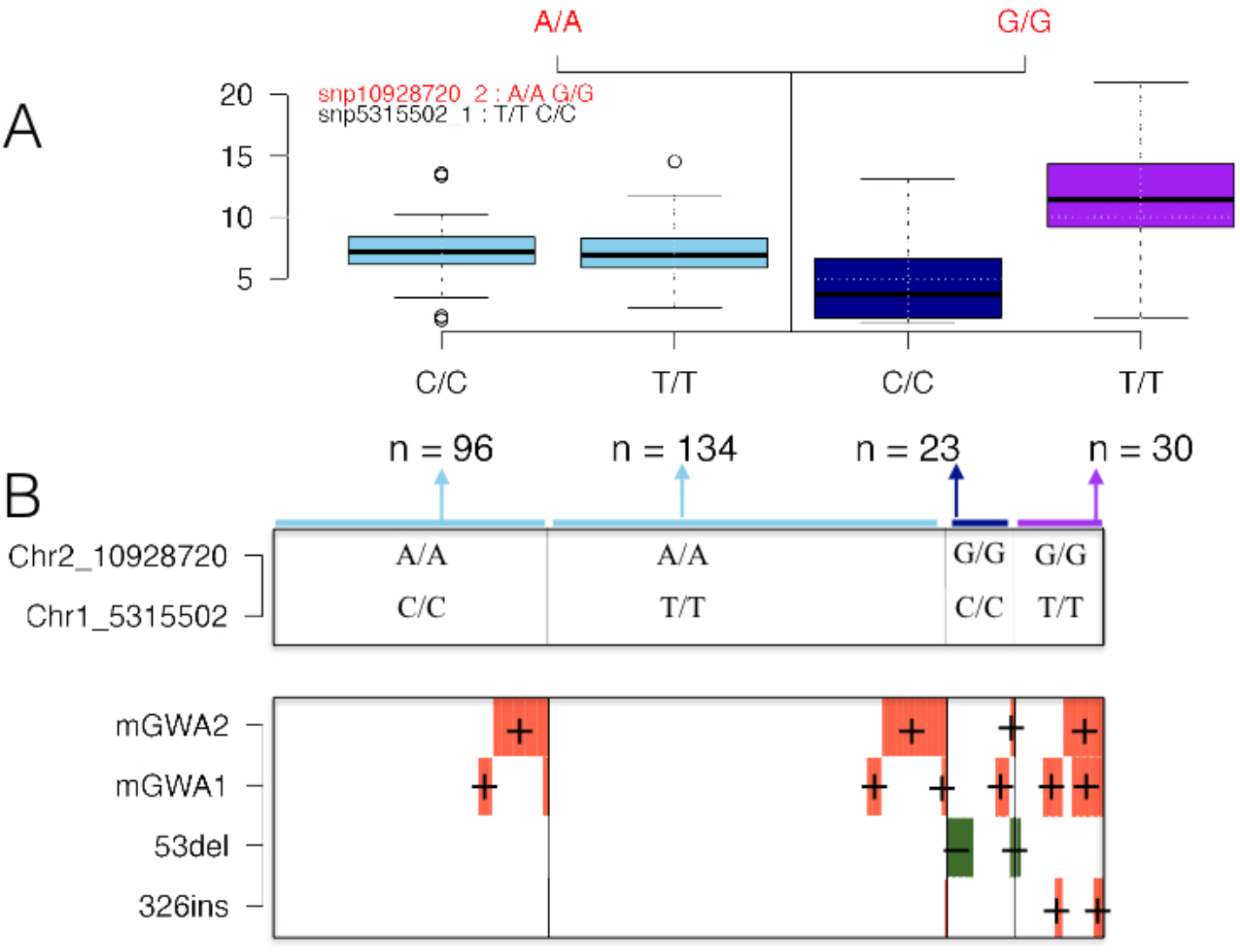
*An illustration of how the high order LD between four polymorphisms affecting the level of molybdenum in the* A. thaliana *leaf (Forsberg et al. 2015), likely explains the significant statistical epistasis detected for a cis-trans predictor pair. **(A)** Boxplots illustrating the phenotypic distribution in the four genotype classes defined by the cis-trans predictor pair with the strongest significant epistatic interaction to the level of molybdenum in the* A. thaliana *leaf. **(B)** Illustration of the connection between the two-locus genotypes of the predictor pair and the minor alleles at the four linked loci associated with this trait on chromosome 2 (Forsberg et al. 2015). The top box in **(B)** illustrates the two-locus genotype for the predictor pair, with the width of each sub-box indicating the number of individuals in each genotype class in the population. In the bottom box in **(B)**, each individual is represented as a column, where green (molybdenum decreasing) and orange (molybdenum increasing) colors indicates that the individual carry the minor alleles at the four loci identified in (Forsberg et al. 2015). mGWAl and mGWA2 are SNP markers associated with the trait and 53del and 326 are structural polymorphisms (Forsberg et al. 2015)*.

This statistical interaction could be due to a true genetic interaction. An alternative explanation is however presented in Figure 5 There, the overlap between the two locus genotypes for the cis-trans predictor pair (Figure 5A) and the alleles at the four loci earlier reported to be associated with leaf molybdenum levels in this region (Forsberg *et al.* 2015) are illustrated. The multi-locus genotypes of the predictor pair tags different combinations of minor alleles at the four loci that were found to either increase (mGWA1, mGWA2, 326ins) or decrease (53del) leaf molybdenum levels in the accessions (Forsberg *et al.* 2015). The statistical epistatic interaction was detected due to the difference in molybdenum levels between accessions carrying the GGCC genotype (low molybdenum) and GGTT (high molybdenum). Figure 5B shows that the accessions with the GGCC genotype have the lowest frequency of the molybdenum increasing allele mGWA2 and the highest frequency of the molybdenum decreasing allele 53del. The accessions with the GGTT genotype instead have the highest frequencies of the molybdenum increasing alleles at mGWA2, mGWA1 and 326ins. The genotypes AACC and AATT, with intermediate molybdenum levels, both have intermediate frequencies of the mGWA1 and mGWA1 increasing alleles and lack the 53del and 326ins alleles. A more parsimonious interpretation of these results is thus that the statistical epistasis at the predictor pair is due to the high order LD between them and the genotypes at the four loci located in the region on chromosome

## Discussion

High order linkage disequilibrium between combinations of genotyped markers, and unobserved functional polymorphisms, can result in significant statistical epistasis in genome wide association analyses. This was earlier illustrated empirically for linked pairs of genotyped predictor SNPs and ungenotyped target polymorphisms in humans by Wood *et al.*(Wood *et al.* 2014). Here, we present a new example from *A. thaliana* where significant statistical epistasis between pairs of predictors is due to the effects at a single loci and that only one of the statistically interacting loci was located near the target. By exploring the prevalence of second order LD in the genome of the 1001-genomes *A. thaliana* collection, we find that although the total amount of high order linkage disequilibrium decreases with increasing population sizes, it is still highly prevalent both within and across chromosomes even in relatively large populations (n = 728). It is was found to be most common when one predictor is in high LD to (and located physically near) the target, but many cases exist where the LD to the individual predictors is very weak but the high order LD is strong. The choice of target and predictor SNPs in this study is arbitrary and therefore it it is difficult to assess how representative they are for the prevalence of high order LD in other populations. However, they do suggest that strong high order LD can be prevalent also in larger populations, indicating that statistical epistasis observed in studies based on reduced representation genotyping (such as SNP-chips) need to be interpreted with caution.

The most prevalent type of high order LD on a genome wide basis is that of cis-trans predictor pairs, but also cis-cis pairs are common regardless of population size. The prevalence of trans-trans pairs is high in smaller populations but decreases rapidly as the population size increases. A possible biological explanation for the observation that cis-cis and cis-trans high order LD pairs is relatively prevalent also at larger population sizes would be that the number of, and variation in, the trans located predictors is sufficiently large on a genome-wide basis to complement any imperfection in the tagging of the functional polymorphism by the cis located predictor. Whereas trans-trans high order LD will always result in falsely associated loci, cis-trans and cis-cis high order LD presents an opportunity to identify true functional loci for the trait. The problem in a real data analysis is that statistical epistasis between a pair of predictors can emerge from true interactions or high order LD within and across chromosomes. However, as the sample sizes increase the risk of detecting pairs of predictors where none is located close to the true functional polymorphism decreases. Before concluding that the detected association is due to two interacting loci, further analyses of the associated pair are however recommended.

Whole-genome sequencing provides unprecedented opportunities to genotype most segregating single nucleotide polymorphisms in the genome. Despite this, it is unlikely that these will be able to tag all functional polymorphisms, such as larger structural variants or multi-allelic functional loci due to tandem repeats. Hence, even though the scenario of reduced representation genotyping with SNP-chips or similar will soon be a technology of the past, association analyses will still be challenged by the need to tag hidden polymorphisms with imperfect markers as illustrated in our analyses of the complex locus affecting molybdenum levels in the *A. thaliana* leaf. In fact, it is not unlikely that the problem with high order LD between SNP predictors and hidden, complex functional loci will remain a major challenge in the future as the increased number of markers generated by sequencing also increases the chance of finding combinations of cis-cis or cis-trans predictors that tag these functional polymorphisms better than any single marker. To evaluate the extent of this problem one will, however, need a more comprehensive dataset than the one studied here including a more complete scoring of all types of non-SNP polymorphisms in the genome with potential effect on traits of interest.

The prevalence of high order LD is likely to be more of a concern in populations of inbred or haploid individuals. These include, for example, inbred lines derived from bi - and multi-parental crosses of plants and animals, as well as populations of wild collected inbred plants (Churchill *et al.* 2004; Valdar *et al.* 2006; Kover *et al.* 2009; Cao *et al.* 2011; Mackay *et al.* 2012). As heterozygotes are not present in these populations, the number of multi locus genotype classes is smaller than in outbred populations, making them attractive for studies of genetic interactions. As a common approach to detect interactions in such populations is to identify pairs of loci displaying significant statistical epistasis, such results need to be interpreted with caution, as the analyzed populations are generally small. If one, or more, of the functional polymorphisms in the genome are unknown and poorly tagged by the genotyped markers, there is a risk that statistical interactions arise from high-order LD between the genotyped markers and the hidden functional polymorphisms. Hence, even though these populations increase the power to map loci displaying statistical epistasis, there is also a risk of falsely concluding that the underlying genetic architecture involves genetic interactions.

## Conclusions

Statistical epistasis detected in genome wide association analyses can result from high order LD between genotyped markers and unobserved functional polymorphisms. This study revealed extensive and strong genome wide high order LD between pairs of markers on a high density 250k SNP-chip and individual markers revealed by whole genome sequencing in the *A. thaliana* 1001-genomes collection. The high prevalence of strong high order LD in this dataset suggests that epistatic variance detected between pairs of markers in association analyses, especially in small inbred populations genotyped for reduced representation sets of markers, need to be interpreted with caution. An empirical example is presented where a pair of markers with significant statistical epistasis in a genome wide association analysis is in high order LD with a complex multi allelic locus with large effects on the analyzed trait. As complex functional loci such as this are unlikely to be captured by individual bi-allelic SNP markers, even if millions of them are scored by whole genome sequencing, it is important to evaluate also other explanations of statistical epistasis than underlying genetic interactions in particular when small populations of inbred individuals are studied.

## Acknowledgements

This study was funded by grants from the Swedish Research Council (2012-4634) and the Swedish Research Council Formas (2013-450) to ÖC.

## Author contributions

ÖC and SF initiated the study. ÖC, SF and YZ designed the project and the statistical analyses; SF and YZ wrote analysis scripts and performed the data analyses. ÖC and YZ summarized the results and wrote the initial version of the manuscript. All authors contributed to the writing of the final version of the manuscript.

## Supplementary Material

Supplementary material is provided in Supplementary Figure 1 and Supplementary Table 1.

